# Novelty detection in early olfactory processing of the honey bee, *Apis mellifera*

**DOI:** 10.1101/2021.10.06.463371

**Authors:** H. Lei, S. Haney, C. Jernigan, X.J. Guo, C.N. Cook, M. Bazhenov, B.H. Smith

## Abstract

Animals are constantly bombarded with stimuli, which presents a fundamental problem of sorting among pervasive uninformative stimuli and novel, possibly meaningful stimuli. We evaluated novelty detection behaviorally in honey bees as they position their antennae differentially in an air stream carrying familiar or novel odors. We then characterized neuronal responses to familiar and novel odors in the first synaptic integration center in the brain – the antennal lobes. We found that the neurons that exhibited stronger initial responses to the odor that was to be familiarized are the same units that later distinguish familiar and novel odors, independently of chemical identities. These units, including both projection neurons and local neurons, showed a decreased response to the familiar odor but an increased response to the novel odor. Our results suggest that the antennal lobe may assign a category of familiarity or novelty to an odor stimulus in addition to its chemical identity code. Therefore, the mechanisms for novelty detection may be present in early sensory processing, either as a result of local synaptic interaction or via feedback from higher brain centers.

## Introduction

Animals are constantly bombarded with stimuli to which they must assign meaning [2, 3]. To assess the importance of a stimulus, information needs to be retained in a limited capacity ‘working memory’ [5-7] for a short period of time. Working memory serves as a preprocessor and an important interface between sensory processing and higher order cognitive functions [8], and the ability to shift the focus of limited attention away from meaningless stimuli and toward potentially meaningful ones is very important. Accordingly, associations between stimuli held in working memory and important events – such as food or threats – can make stimuli more salient [9, 10], whereas the lack of any salient association reduces an animal’s attention to inconsequential stimuli. The latter occurs in Latent Inhibition [11-13], where animals habituate to repeated stimulation by ‘familiar’ stimuli. The process of familiarization can also lead animals to pay more attention to ‘novel’ stimuli being experienced for the first time. Although novelty detection can be measured by heightened behavioral responsiveness to a novel stimulus, it is not yet well documented where or how novelty detection might be implemented in neural circuits in the nervous systems.

The primary olfactory center, the antennal lobe (AL) in insects and olfactory bulb in vertebrates, is well situated to perform critical odor preprocessing and to categorize olfactory inputs on glomerular activity maps for odors [14, 15]. Within each glomerulus, extensive synapsing occurs between sensory neurons and interneurons, and between the interneurons themselves. Both associative and non-associative plasticity at different synapses in the AL [16-20] bias odor representations to allow animals to focus a limited attention capacity on salient, predictive stimuli [11, 19, 21, 22]. This plasticity is driven, in part at least, by release of octopamine from Ventral Unpaired Medial neurons in the subesophageal ganglion of the brain when the honey bee feeds on sucrose [23, 24]. Attention is also modulated by novelty and familiarity, which are experience-dependent qualities. The same odor stimulus can be novel or familiar to an animal depending on the animal’s previous experience. There are no specialized novelty or familiarity sensing neurons on the antennae or known specialized glomeruli in the AL to mark odors being novel or familiar. Therefore, novelty and familiarity are new qualities that are created within the neural networks in the AL or elsewhere in the brain that provide a substrate for modifying working memory.

In the fruit fly, *Drosophila melanogaster*, it has been shown that synaptic plasticity mediated by GABAergic mechanisms in AL circuits underlies a stereotypical response to familiar odors in an odor-specific manner [25], but it is unclear if and how the same circuit motif affects responses to novel odors. On the other hand, the mushroom bodies, which are the third-order synaptic centers along the olfactory pathway and important centers for learning and memory, generate distinct response patterns to both familiar and novel odors [26]. In particular, one type of mushroom body output neuron, namely α’3 MBONs [27], decrease response magnitude to familiar odors over repeated stimulations but respond strongly to novel odors. The novelty detection requires participation from a dopaminergic neuron innervating the same structure [26]. However, it is not known whether the neural code for novelty and familiarity is generated only in the mushroom bodies, or whether it can be reflected in earlier circuits, such as the AL, through intrinsic synaptic modifications or via feedback from the mushroom body.

Here, we show in the honey bee that AL local and projection neurons can be potentiated to novel odors and adapted to familiar odors after unreinforced exposure - familiarization - to an odor. This provides a neural coding scheme to categorize odors as being novel or familiar in addition to the glomerular coding patterns for chemical identities.

## Results

### Different antenna movement to stimulation by novel and familiar odors

We began by asking if there are observable behaviors by which honey bees could exhibit novelty detection. We used a familiarization protocol called Latent Inhibition [13], which has been established in proboscis extension response (PER) conditioning of honey bees [12, 28] to familiarize subjects to one of two odors (Fig. 1A). Previous PER studies have shown that 20 to 40 presentations of an odor without reinforcement slows honey bees’ ability to associate this now familiar odor with reinforcement relative to a different, novel odor [12]. We now evaluated how familiarization might be exhibited in honeybee antennal movements to odors. The bees were restrained in harnesses that are commonly used in procedures for studying classical and operant conditioning in PER [29], and which allow monitoring of antennal movements. Restrained honey bees actively scan the air flow around them with their antennae to sample headspace volatiles [30]. The patterns of these movements can be modified by appetitive conditioning [31, 32]. Using video tracking [30], we recorded, frame-by-frame, the angular positions of each of the antennae in a 180° space on each side of the head. We recorded positions continuously in 4 sec blocks before, during, and after exposure to two odorants, 1-hexanol (hex) and 2-octanone (oct) (Fig.1A).

**Figure 1.**
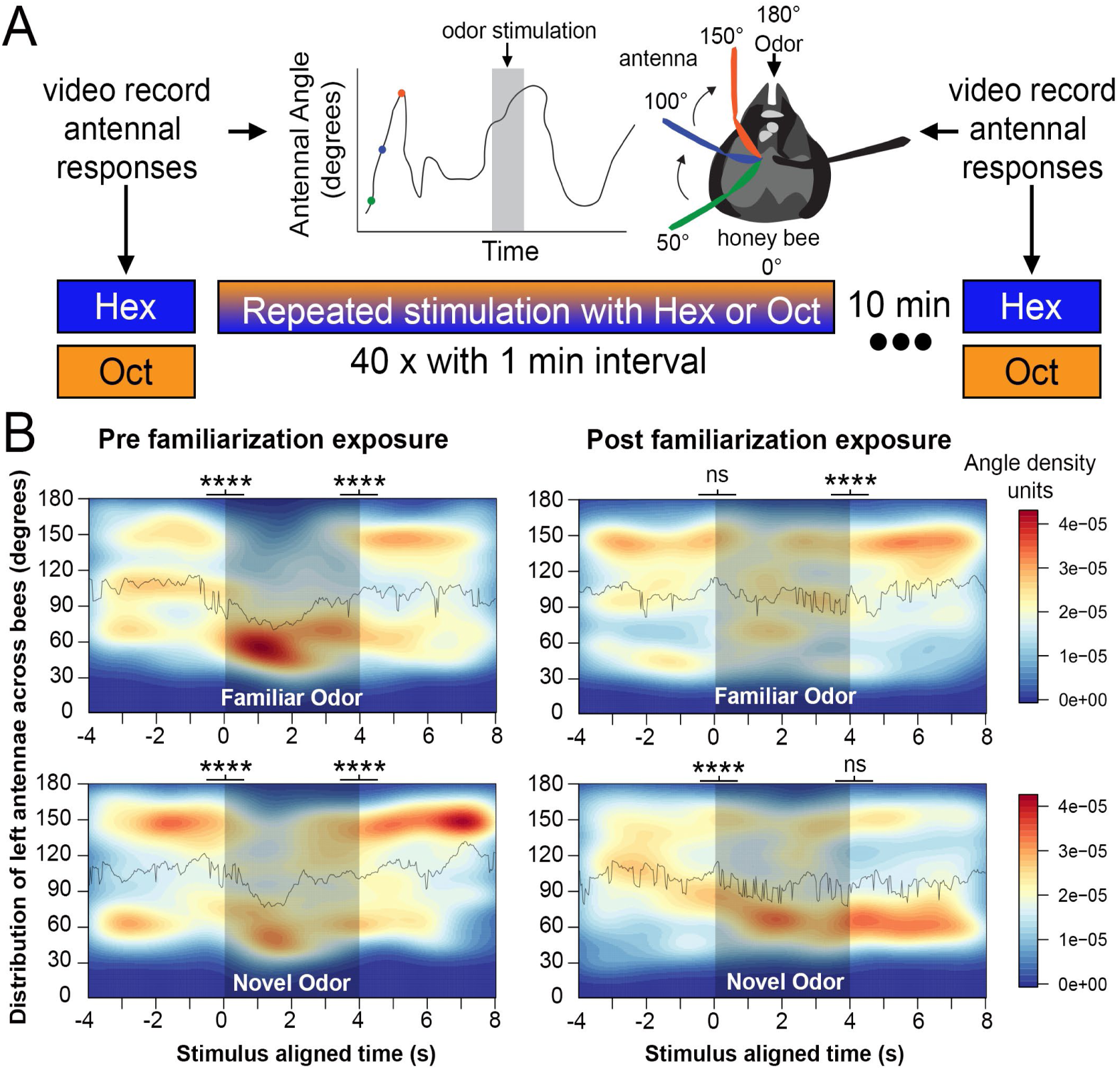
Left antenna responses to novel and familiar odors. **A**. The familiarization protocol using either Hexanol or Octanone as the repeatedly presented unreinforced odor stimulus. Tests were performed with each odor prior to and after familiarization. The headspace is divided into a clockwise circular range of 180 degrees for each antenna. **B**. The average distribution densities of left antennae’s angular positions in response to odors before the familiarization process (left column plots) and 10 min after the familiarization process (right column plots). Plots are aligned by odor onset over 12 s, which includes 4 sec segments before, during and after odor delivery (360 frames; N=24 animals, 13 Hex exposed, 11 Oct exposed). In both columns, the top plot shows responses to the familiar odor and the bottom plot shows responses to the novel odor. The solid line in each plot represents the mean angle of the left antenna across all 24 bees for each respective odor. The shaded region indicates the period during odor stimulus presentation. The colored scale to the right of the plots shows the standardized Kernel density scale across all plots. For statistical comparisons angular position was normalized by calculating the difference in the left antenna’s angular position at all time points from the mean antennal angle (when no odor present) across the three time-sections for each bee and odorant presentation. Statistical analyses: two-way ANOVA, difference measure∼ time period * odor, df=2, n=24, Tukey HSD post-hoc comparison, asterisks indicate statistical significance, **** p<0.00001, ns indicates not significant.

Prior to becoming familiar with the odors, honey bees responded to both odors similarly (Fig.1B). In the period before the odor is presented, honey bees sweep their antennae forward and backward and the antennal positions reflect a fairly equal probability of being in typical areas of the space, i.e. equal forward, central and backward distribution. The backward position is comparable to a lifted antenna position in bee’s natural posture when walking or flying. During the 4 sec of odor presentation, the antennae are mostly held backward, and during the 4 sec just after the odors are switched off, the antennae return to a more typical distribution across the space (Fig.1B). Transitions from before to during and from during to after were significant in all cases. The asterisks at the border between different time sections in Fig.1B indicate that the difference in antenna’s position from the mean antennal angle (when no odor was present) is statistically significant between the time blocks (two-way ANOVA, n=24, Tukey HSD post-hoc comparison, **** p<0.00001).

The pattern of significance prior to familiarization shows that initial responses to the soon to be novel and familiar odors were similar. In contrast, after the familiarization process the same transitions differed for the familiar and novel odors (Fig.1B). The now familiar odor no longer elicited such sharp transitions across the three time-segments. However, the ‘novel’ odor showed the expected transition from before to during the odor presentation. Importantly, the antennae maintained the backward position even after the novel odor was terminated. Movements of the right antennae also showed similar patterns to the novel odor (the pattern changed when the novel odor was presented and did not change when it was terminated), although the diminished response to the familiar odor was less pronounced (Supplemental Fig 1). These backward movements, and in particular the persistence in those movements to a novel odor, imply that they are involved in odor detection, and we interpret them as indicating the degree of novelty or surprise in a new odor.

**Supplemental figure 1.**
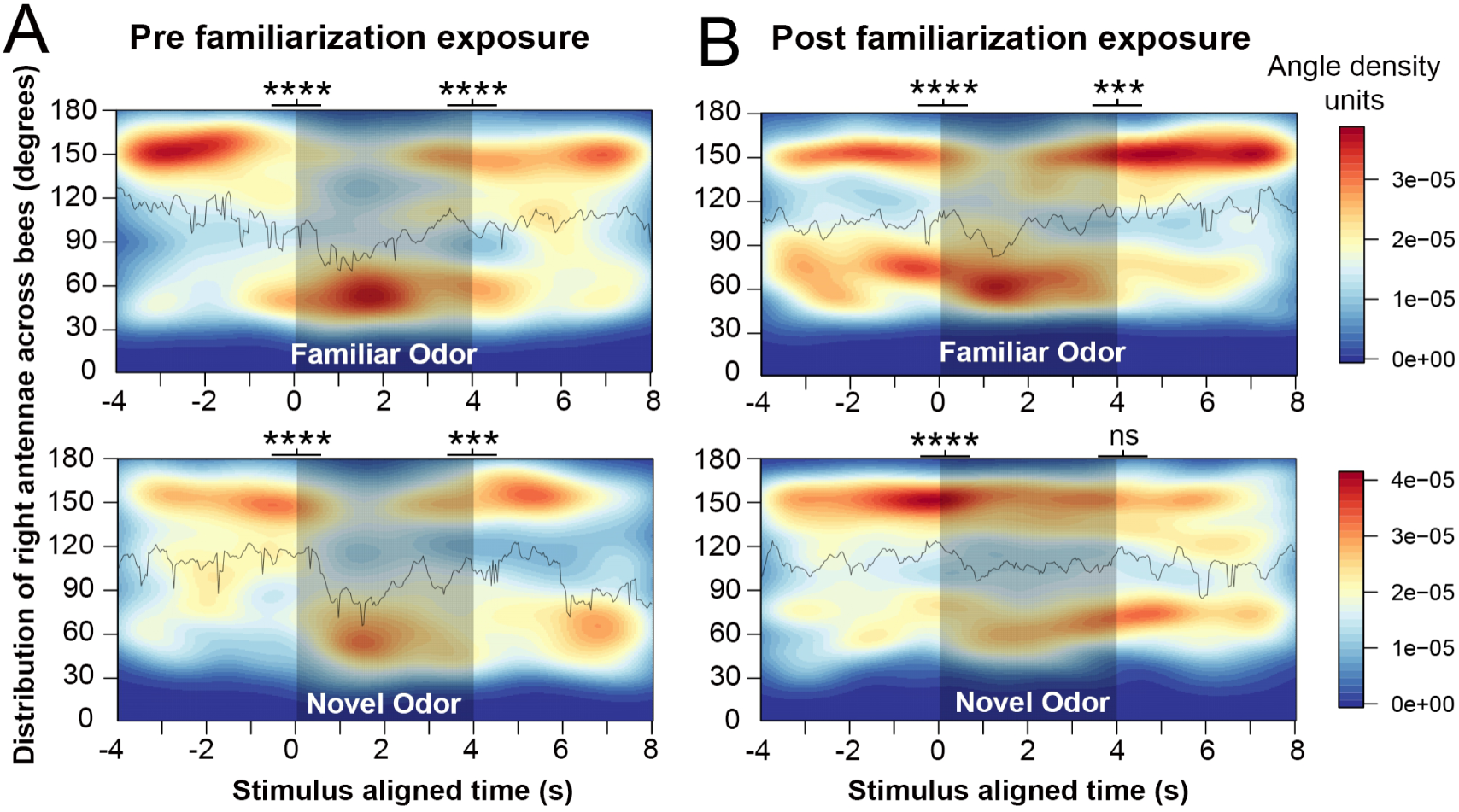
Right antenna movement to novel and familiar odors. Angular positions are defined as in figure 1. **A**. The average distribution densities of right antennae’s angular positions in response to odors before the familiarization process (left column plots) and 10 min after the familiarization process (right column plots) aligned by odor onset over 12 s Statistical analyses: two-way ANOVA, difference measure∼ time period * odor, df=2, n=24, Tukey HSD post-hoc comparison, asterisks indicate statistical significance, **** p<0.00001, ns indicates not significant.

### Familiarity and novelty effects on PNs and LNs

We then sought neural correlates of the novelty-dependent behavior in the AL. In the insect AL [33], all of the olfactory receptor neurons in an antenna that express the same receptor project axons to the same area (glomerulus) of the AL, where they synapse with PNs that relay activity to higher brain centers such as the Mushroom Body. Different types of Local Interneurons (LN), which are mostly GABAergic, interconnect glomeruli and set up local processing in the AL. We recorded two hundred seventeen units from the left AL of 19 bees before and after inducing familiarity using the same familiarization protocol as above (Fig. 1A) [28]. These extracellularly recorded units were classified into probable PNs (N=87) and LNs (N=115) based on the statistical properties of their spiking patterns. Specifically, following methods outlined by Meyer and colleagues [1], we assessed the variation of the inter-spike interval within a trial, the variation of spike count across trials, the change in firing rate due to the stimulus, and the lag-time to response (see Methods, Fig. 2A, B). These features were sufficient to classify two different categories of units whose properties agreed with morphologically identified PNs and LNs (Fig. 2C-E and see Meyer et al. 2013). Some units (N=15) were not able to be classified because of low spike counts.

**Figure 2.**
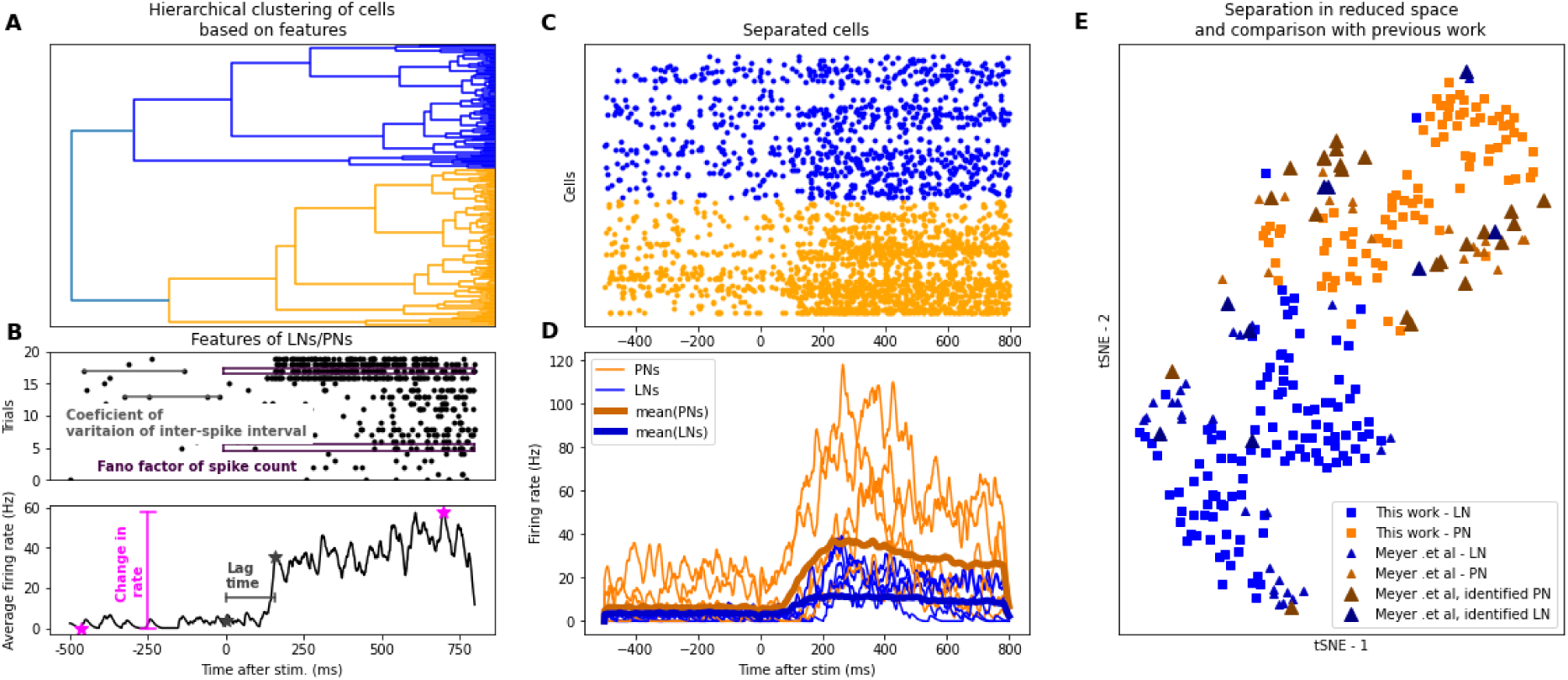
Separation of PNs and LNs using a statistical procedure outlined in [1]. **A**. Separation of LNs and PNs using hierarchical clustering (see Methods) into LNs (blue) and PNs (orange). **B**. Explanation of cell-based features used for clustering (see Methods) based on spike times (top) and response rate (bottom). **C**. Spike rasters and, **D**. firing rate time courses of separated cells. **E**. Cell-based features visualized in reduced space using tSNE (t-distributed Stochastic Neighbor Embedding) [4].

We quantified a neuron’s ‘net response’ to odor by calculating a ratio of the amount of stimulus-evoked response relative to the spiking activities for an equivalent period immediately prior to stimulation (see Methods). This metric allows a closer comparison to previous experiments using calcium imaging, where ΔF/F_0_ was calculated [17, 34]. We compared the averaged responses of both PNs and LNs separately to the familiar and novel odors before and after familiarization. We first pooled the stimuli into familiar or novel categories irrespective of the specific odors (Hex or Oct) used in the familiarization process. As a result of the familiarization process, many PNs (Fig.3 A1, B1) and LN’s (Fig.3 A2, B2) showed stronger responses to the novel odor. On average, for both PNs and LNs there was a reduction in response to the familiar odor and an increase in response to the novel odor after familiarization relative to before (Fig. 3 C1 C2). To test the significance of this pattern, we performed 2-way ANOVA using the log normalized pre- and post-familiarization responses as one factor and responses to familiar and novel odors as the second factor. For both PNs and LNs the interaction between the two main effects was significant (PNs: F = 4.06, p < 0.05; LNs: F = 4.7, p < 0.05). That is, the decrease in response to the familiar odor between pre- and post-familiarization significantly contrasts with an increase in response to the novel odor. This pattern was largely confirmed with post-hoc t-tests (Fig 3). For both PNs and LNs the increase in response to the novel odor was significant. The decrease in response to the PN was marginally significant for the PNs and not significant for the LN. These data suggest that novelty potentiates responses in PNs and LNs, and familiarity may decrease responses to PNs.

**Figure 3.**
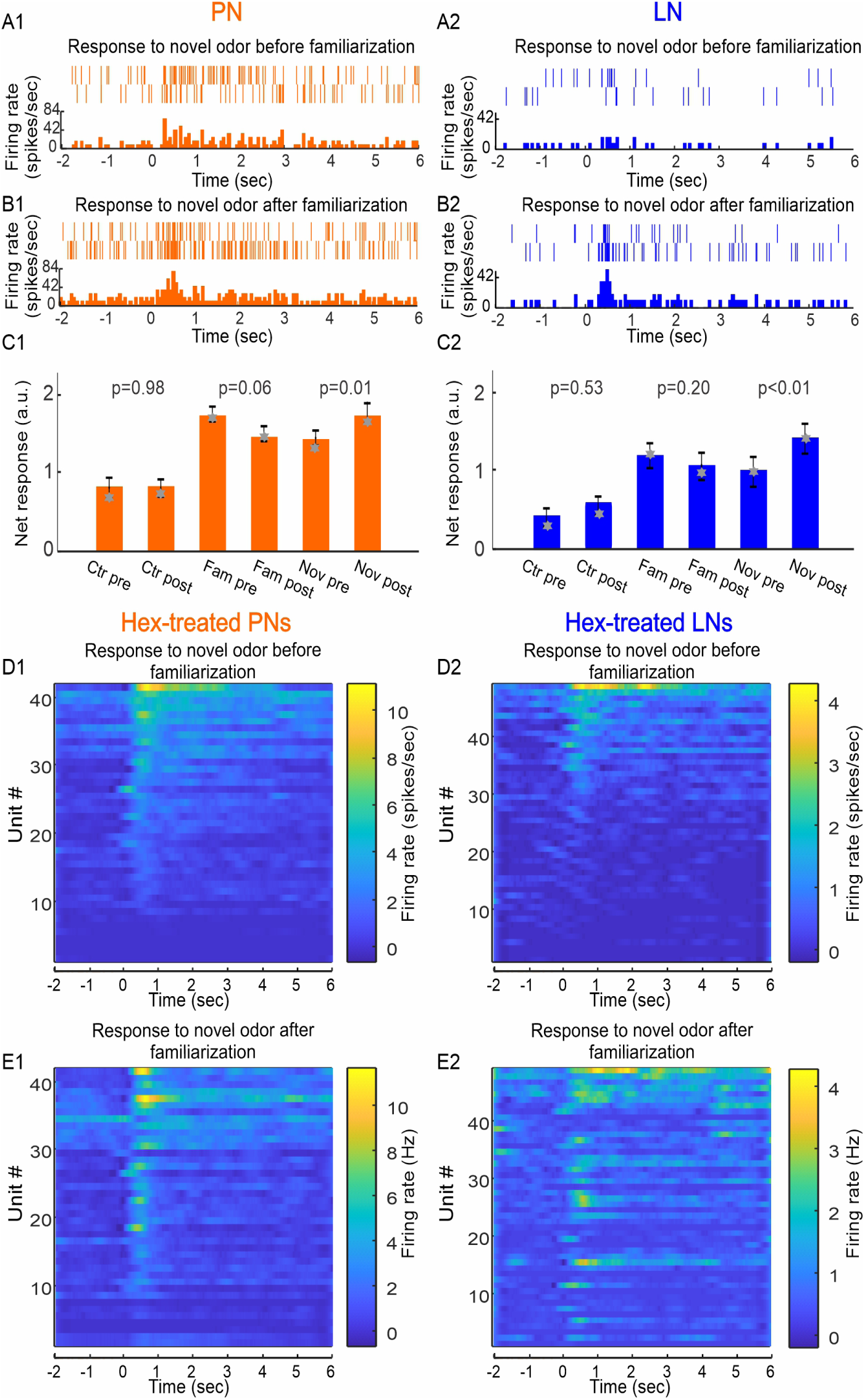
Novelty and familiarity is evident in both PNs and LNs. Familiarization potentiates the responses of PNs (A1, B1) and LNs (A2, B2) to novel odor (Hex), evident by the raster plots (top of each panel) and peristimulus time histograms (PSTHs) (bottom of each panel). The odor was delivered at time zero, lasting for 4 sec. The mean response of all PNs (orange bar graph, C1; log transformed) shows a marginally significant decrease to the familiar odor from pre- to post-familiarization (middle two bars, mean ± S.E., p=0.06, n=87, paired-t test), and a significant increase to the novel odor (right two bars, mean ± S.E., p=0.01, n=87, paired-t test); there is no significant change in response to the solvent control (left two bars, mean ± S.E., p=0.98, n=87, paired-t test). In LNs (C2), the familiarity effect is not significant (middle two bars, log transformed; mean ± S.E., p=0.2, n=115, paired-t test) but the novelty effect is significant (right two bars, mean ± S.E., p<0.01, n=115, paired-t test). No significance was detected to solvent control (left two bars, mean ± S.E., p=0.53, n=115, paired-t test). The grey stars in C1 and C2 marks the median value, which was used to establish rough equivalence to the mean, as would be the case for a normal distribution needed for parametric tests. The pseudo-colored PSTHs of all Hex-familiarized PNs (F1, G1), ranked low (bottom) to high (top) according to firing rate pre-familiarization (F1), show clear variation across units in the novelty effect before and after familiarization. The novelty effect appears to be even more variable among the Hex-familiarized LNs (F2, G2).

To further visualize the response patterns in PNs and LNs, we plotted the peristimulus time histograms generated by the Hex-familiarized PNs and LNs (Fig.3 D1, E1, D2, E2). Comparison of PN (D1 and E1) and LN (D2 and E2) responses pre- and post-familiarization reveal changes in some units in responsiveness to the novel odor, which is the basis for the average differences in responses (C1 and C2). However, there is clearly variation among the neurons in their capacity to differentiate familiar from novel odors, thus necessitating analyses based on further sub-classifications among PNs and LNs.

### Novelty and familiarity effect on response-based subtypes of PNs and LNs

A subset of PNs and LNs showed biased responses to Hex or Oct prior to familiarization (Fig.4 A1, A2). We therefore classified each PN and LN as either familiar-odor biased or novel-odor biased, based on which odor it showed a stronger response to and on which odor was eventually used for familiarization. We then evaluated novelty detection on these different subsets by calculating the change in response due to familiarization for the novel and familiar odors in either Hex-familiarized or Oct-familiarized PNs and LNs (Fig.4 B1, B2). For Hex-familiarized PNs, the novel odor was Oct and the familiar odor would be Hex, and visa-versa. In familiar-biased PNs, there were more neurons showing stronger response to the novel odor after familiarization, depicted by the positive median values of the 2^nd^ and 4^th^ boxes in Fig.4 B1. In contrast, these PNs showed decreased response to the familiar odors after familiarization, depicted by the negative median values of the 1^st^ and 3^rd^ boxes. A similar pattern, i.e. increases in responses to novel odor and decreases in responses to familiar odor, was also observed in the familiar-odor biased LNs (Fig.4 B2).

**Figure 4.**
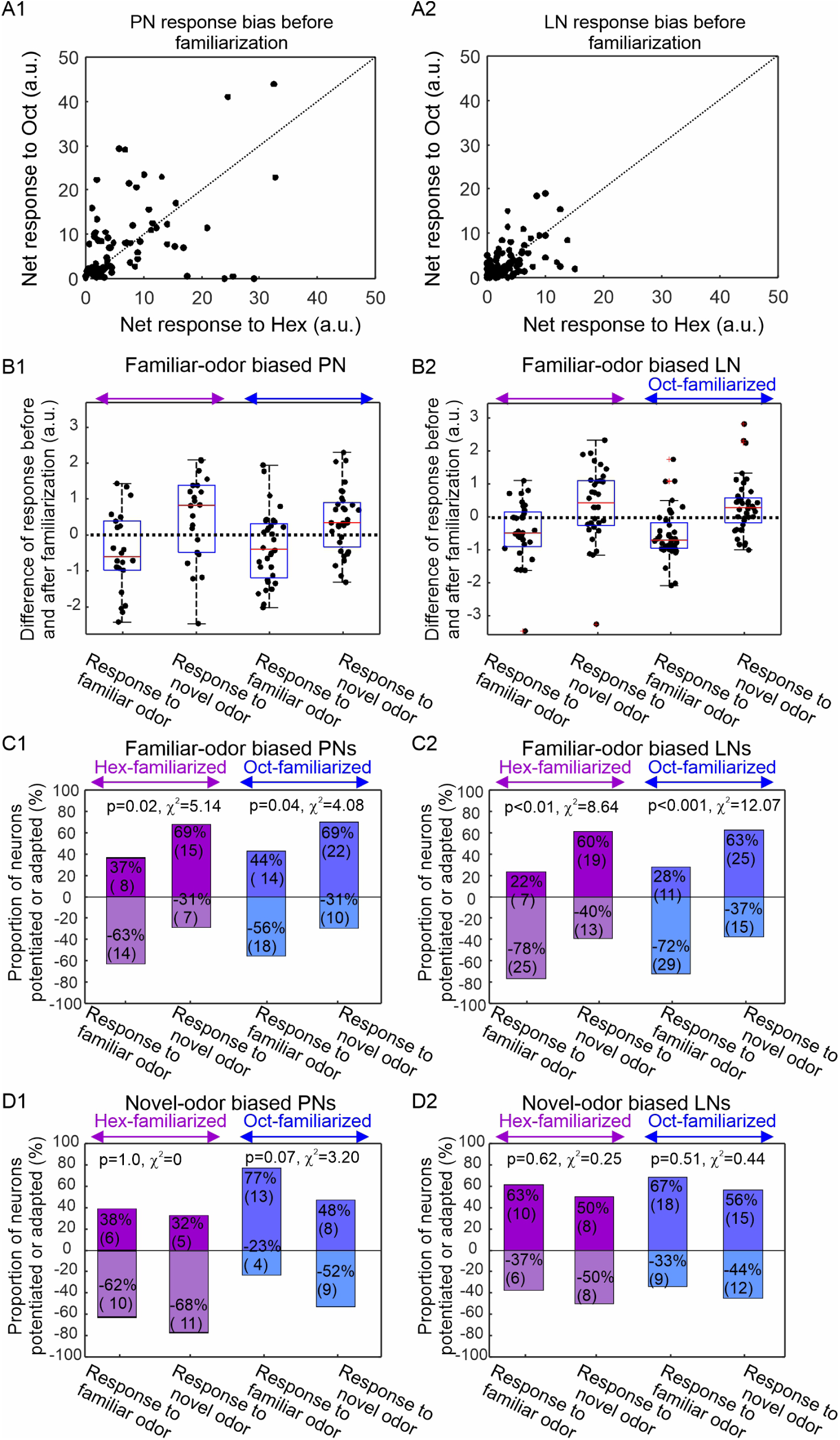
Familiar-odor-biased PNs and LNs are novelty and familiarity detectors. PNs and LNs display initial response biases to the two test odors Hex and Oct prior to familiarization (A1, A2). Some neurons respond more strongly to Oct than to Hex, i.e., data points above the diagonal line; some neurons respond more strongly to Hex than to Oct, i.e., data points below the diagonal line. Additionally, PNs are generally more responsive than LNs, evident from the wider distribution of data points along both X and Y axis. The Oct-biased units are above the diagonal line and the Hex-biased units are below the diagonal line. Then they are familiarized with Oct or Hex. Only the familiar-odor biased neurons showed consistent familiarity and novelty effects. In familiar-odor biased PNs (B1), whether it is Hex-familiarized or Oct-familiarized, most of the neurons decreased their responses to the familiar odor after the familiarization process, as revealed by the negative median values in the box plots (1^st^ and 3^rd^ boxes). In contrast, most of the neurons increased their responses to novel odor after the familiarization process – namely novelty effect, as shown by the positive median values in the box plots (2^nd^ and 4^th^ box). The familiarity and novelty effects are also evident in the familiar-odor biased LNs (B2). In Hex-familiarized familiar-odor biased PNs, 63% (n=14) of the neurons exhibited familiarity effect and 69% (n=15) of the neurons exhibited novelty effect; this distribution pattern is significantly different from random process (p=0.02, McNemar test) (purple boxes, C1). In Oct-familiarized familiar-odor biased PNs, also seen were significantly higher percentage of neurons adapted by familiar odor (56%, n=18) and potentiated by novel odor (69%, n=22) (blue boxes, C1, p=0.04, McNemar test). Similar phenomena were also observed among the familiar-odor biased LNs (C2). In novel-odor biased PNs and LNs, the familiar and novel odors did not produce contrasting changes after the familiarization process; none of the comparing pairs showed significant familiarity and novelty effects (D1, D2, McNemar test).

This pattern is statistically significant in both PNs (Fig. 4 C1; McNemar test, p=0.02 for Hex-familiarized group; p=0.04 for Oct-familiarized group) and LNs (Fig. 4 C2; McNemar test, p<0.01 for Hex-familiarized group; p<0.001 for Oct-familiarized group). These results indicate that neurons with a strong initial response to the familiarization odor showed the most dramatic changes both to the familiar and novel odors during the familiarization process (Fig.4 B1, B2, C1, C2). In contrast to the familiar-odor biased neurons, neither novel-odor biased PNs (Fig.4 D1) nor novel-odor biased LNs (Fig.4 D2) showed significant changes induced by familiarization (PNs: McNemar test, Hex-treated p=1.0, and Oct-treated p=0.07; LNs: McNemar test, Hex-treated p=0.62, and Oct-treated p=0.51).

These results imply that familiar and novel odors should become more distinguishable after the familiarization process among neurons for which Hex and Oct elicit strong responses. This prediction was verified by comparing the spatial separation between these two odors before and after familiarization in a coding space constructed from the first three principal components of PN population responses to familiar and novel odors (Fig.5A). Moreover, the largest separation occurred at the peak response after odor stimulation, as shown by the Euclidean distance curves calculated from the difference of PN population responses to familiar and novel odors before and after familiarization (Fig. 5B).

**Figure 5.**
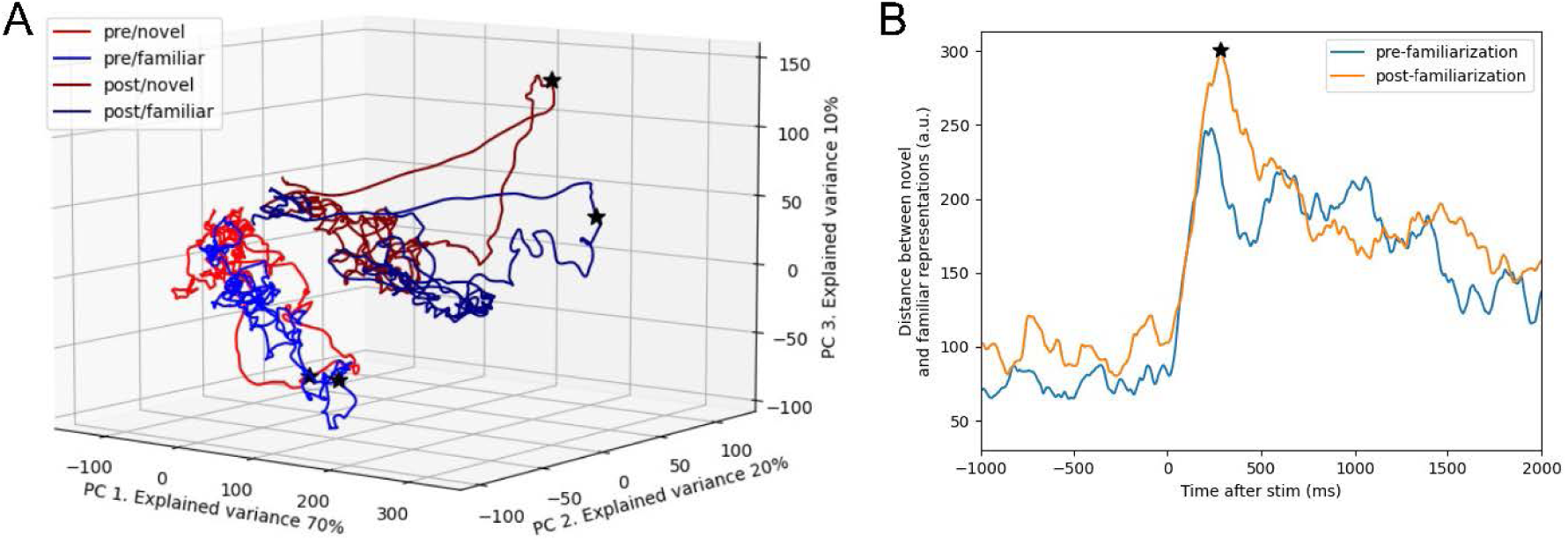
Familiarization enlarges the differences between familiar- and novel-odor representations in AL. **A**. Principal components of firing rate of PN responses to familiar- and novel-odor before and after the familiarization process (pooled across hex-familiarized and oct-familiarized PNs). Black stars mark the time points where the peak response was. After familiarization, the novel odor evoked a response that showed distinct prominence at peak response relative to all other presentation types. **B**. Familiarization produced increases in the distance between the novel and familiar PN response (pooled across hex-familiarized and oct-familiarized PNs). Black star occurs at the same time as in A. Stimulus onset was at 2s.

## Discussion

Novelty perception is a context-dependent phenomenon. Behaviorally, animals tend to pay more attention to novel stimuli than to familiar ones. Using antennal tracking we have developed a method for identifying novelty detection in harnessed bees. We also found evidence of novelty detection in the primary olfactory neuropil, the AL. Overall, novelty detection in the AL appears to expressed in the group of neurons biased to the familiar odor.

### Antenna movement affected by familiar and novel odors

As a pair of prominent sensory organs, insect antennae move in specific ways upon stimulation. Cockroaches modify antenna movements in opposite ways in response to attractive and aversive odors [35]. Honey bees move their antennae backwards to odor stimulation before conditioning, and then forward after odor has been conditioned to sucrose reward [31]. In our study we also found that honey bees hold their antennae at a backward position when stimulated by odor. We also show that this movement is modified after odors take on meaning. Specifically, that movement is attenuated after repeated exposure – familiarization – to an odor, such that its ‘meaning’ is nothing important. In that context, when presented with a novel odor, which may or may not mean something important, honey bees extend the time they hold the antenna in a backward position well beyond termination of the odor. It is possible that this novelty response, and specific odor-driven movements of the antennae in general, affects sensing. After a bee has been conditioned to associate an odor with sucrose reinforcement, it responds to the odor by moving its antennae to the forward point where the bee has received sucrose-water droplets in the past [31]. Movements in this context could therefore enhance detection of both sucrose and odor analogous to how bees move their antennae to inspect flowers [36]. On the other hand, bees may move their antennae in different ways upon stimulation with a novel or familiar odor, which in turn change how the odor is perceived. That is, antennal movements described in our study could be considered as ‘active sensing’ [37, 38], meaning the sensor’s behavior is altered by the bearer in order to enhance ‘meaningful’ signal detection. This strategy could be important, especially when the odor signal is scarce. Odor molecules are carried by turbulent airstreams that shear the clouds of molecules into thin filaments [39]. These filaments pass over the antenna for brief (milliseconds) periods. The movements we describe here could enhance detection of these sparse filaments, thus enhancing the detectability and/or discriminability of odors in an airstream. It could therefore be that the movements we describe here perform an analogous function to increased sniffing in mammals towards novel odors [38, 40].

### Expanded coding dimensions for novelty and familiarity

The one-to-one anatomical relationship between glomeruli in the AL and receptors on antennae suggests that one main function of the AL is to encode the chemical identities of odor stimuli [34, 41, 42]. Odor molecules activate a range of receptor types depending on the molecular identities and binding properties of the receptors [43]. Axons projecting from the olfactory receptor neuron to the AL in turn activate their cognate glomeruli in the AL. The spatiotemporal activity patterns across glomeruli in the AL therefore reflects the response spectrum to different odors [34, 44].

We have identified an important means for representing novelty and familiarity in the early olfactory processing circuits in the AL, which is overlaid on the glomerular map for odor identity. Our data suggest that every glomerulus can participate in the encoding of familiarity and novelty by responding to familiar and novel odors in distinct patterns. These neurons show decreased responses to familiar odors and increased responses to novel odors (Fig.4). We observed the same response patterns irrespective of odor chemical identity, indicating that this phenomenon has the ability to encode a higher-dimensional feature (i.e. familiarity and novelty) in addition to the odor’s chemical identity.

Although PNs and LNs in every glomerulus can encode familiarity and novelty, the odors that can produce familiarity-associated reduction are most likely the odors that the glomerulus is tuned to. This finding is consistent with a *Drosophila* study where the authors identified a glutaminergic mechanism underlying glomerulus-specific habituation [25]. In Drosophila AL, potentiated GABAergic LNs are necessary and sufficient to reduce PN’s response to previously exposed odors. Moreover, these LNs co-release GABA and glutamate. The odor-specificity of habituation is achieved through the activation of NMDA receptors on the PNs. In the current study, we found that PNs and LNs show reduced response after the familiarization process to the odor that they are biased towards initially (Fig.4). This finding is consistent with the odor-specific habituation model in *Drosophila*. Additionally, our study also revealed that these biased PNs and LNs increase response to novel odor – a result that was not reported in the *Drosophila* model. Although we have only tested one novel odor per animal, we infer that other novel odors would produce similar novelty effect because the odors we selected do not have any specific correlation with the familiar odor. In other words, a sustained exposure to the odor that the glomerulus is most sensitive to creates a representation of familiarity by weakened response to this odor; but the same glomerulus may be able to generate a representation of novelty to many or all other non-selective odors. As such, the original glomerular output that is consisted of a whole spectrum of responses evoked by a range of odors is re-organized to reflect previous experience. After familiarization, the most effective odors become less effective and vice versa. Therefore, the adjusted contrast can represent familiarity and novelty. A strengthened response is associated with novel odor, and weakened response with familiar odor, irrespective of molecular identities.

### Novelty encoding in AL and MB

The enhanced sensory patterns driven by novel odors should be integrated into higher order processing in the mushrooms bodies and possibly also in the lateral protocerebrum [45-47]. Recently, a means for detecting novel odors was identified in the fruit fly mushroom bodies [26]. An identified dopaminergic neuron is responsible for the rapid transition from a familiar to a novel state in a group of mushroom body output neurons (MBONs). In that circuit, the novelty detection is not selective, meaning any new odor can release the MBONs from the suppressive state.

There are three hypothetical frameworks for placing our results in the context of the fruit fly results. First, novelty detection in the AL, which we propose here, could be a different mechanism from the dopaminergic mechanism in the mushroom bodies, and it could *augment* the novelty-coding in the mushroom body as the contrast between response to familiar and novel odor in the AL is projected to the mushroom body circuits. Second, novelty detection in the AL of the honey bee might not arise from properties intrinsic to the circuitry of the AL. Instead, the novelty effects we detected might have arisen in the MB and transmitted via feedback neurons to the AL. Such a circuit has been identified in the fruit fly [48] and in anatomical analyses of the honey bee brain [47]. Third, novelty detection may have been pushed from the MB to the AL because of important differences in the roles of different structures in the honey bee and fruit fly brains [49], just as species-specific differences affect homologous areas of mammalian brains [50]. The fruit fly mushroom bodies contain approximately 4,000 intrinsic neurons, and the brain itself contains approximately 135,000 neurons [51]. Thus, the mushroom bodies make up about 3% of the total brain neurons. Inputs to the fruit fly mushroom body are predominantly from the AL and carry olfactory information. In the honey bee, the mushroom bodies contain approximately 340,000 intrinsic neurons, which comprises circa 34% of a brain with just under 10^6^ neurons [49]. The relatively larger size reflects the highly multimodal nature of the inputs from visual, mechanosensory, taste and olfactory processing centers in the honey bee brain [49]. Given the focus on multisensory information processing in the honey bee mushroom bodies, it could be that novelty detection has been pushed earlier in the sensory processing stream, into the AL, which would mean that the mechanisms we describe are independent phylogenetic additions to olfactory processing in honey bees.

### Plasticity as an important component in the AL circuitry

Biogenic amines such as serotonin, dopamine, tyramine and octopamine play important neuromodulatory roles in the honeybee antennal lobe [52]. For example, octopamine and tyramine are released by a cluster of VUM neurons that have cell bodies in the subesophageal ganglion and project processes to most or all glomeruli of the antennal lobe [53]. Those processes arborize in the outer core of glomeruli, where axons from olfactory receptor neurons interact with processes from LNs and PNs [52, 54, 55]. More specifically, octopamine receptors colocalize with inhibitory neurons and tyramine receptors colocalize with axon terminals form receptor neurons, suggesting that these two neuromodulators may play different modulatory roles. Perhaps, similar to the corelease of GABA and glutamate that mediates glomerulus-specific habituation to familiar odors in *Drosophila* AL [25], the corelease of octopamine and tyramine may mediate glomerulus-specific potentiation to novel odors in honey bee AL.

Previous studies using calcium imaging in the honey bee AL have identified both associative and non-associative plasticity. Differential conditioning of two odors [17], or two clusters of odors [56], have shown that the representations for those odors become significantly separated. Computational modeling on the AL circuitry has shown how this increased separation could arise by octopamine actions on inhibitory LNs [16], which has been shown anatomically [52]. Furthermore, experiments using a similar familiarization protocol to the one used here found that familiarization caused a shift in the representation of odor mixtures, i.e. 50-50 mixtures of a novel and familiar chemical components [28]. Specifically, the response to odor mixtures became more similar to the novel odor and less similar to the familiar odor response. However, the biogenic amines underlying this plasticity, or the novelty effect, is not known. Plastic facilitation at the LN-PN and LN-LN synapses could account for this shift in odor representation.

### Summary

We have identified a mechanism by which the honey bee AL distinguishes familiar-from novel-odors independent of their chemical identities. More work now needs to be done to determine how it is implemented, how long it persists, and how it functions in the natural environments of the honey bee. Our results, together with other studies of plasticity in the AL [28, 56], expands the AL coding dimensions to include higher-order features such as olfactory familiarity and novelty. This new coding scheme in the AL, along with the novelty detection at higher synaptic centers such as MB, likely shift attention to novel odors, which may offer behavioral benefits to the animal when linked to active sensing [37, 38].

## Materials and Methods

### Animal preparation

Non-pollen forager honey bees (*Apis mellifera* L.) were collected in the morning at the entrance of a hive maintained outdoor in natural lighting conditions, briefly cooled and restrained in individual harnesses. For behavioral experiments, bees were restrained with tape and a small amount of dental wax behind the head to fix their head in place. After recovering from cooling, bees were then fed 30 % sucrose (w/w) until satiation and allowed to adjust to harnesses for approximately 1 hour in ambient conditions.

For electrophysiological experiments, bees were restrained in harnesses with dental wax around head. After recovering from cooling, bees were fed 2 μl of a 1.0 mol l^−1^ sucrose solution and allowed to remain undisturbed for in a moisturized box until experiments took place. Before recording, the antennae were immobilized with eicosane gently applied to the base of scape and the joint between flagellum and pedicel. A rectangular window was cut open on the cuticle between the compound eyes to expose the antennal lobes. To minimize the muscular and hemolymph movement in the head capsule, the base of proboscis underneath the frontal cuticle plate was severed and the abdomen was removed. A continuous saline flow through the head capsule was supplied during the entire time course of recording.

### Electrophysiological recordings and experiment protocol

Extracellular recordings were performed in the left antennal lobe with a 16-channel probe (NeuroNexus, Ann Arbor, MI). Spike waveforms were digitized with a RZ2 system at a sampling rate of 20 KHz (Tucker-Davis Technologies, Alachua, FL). A total of 217 units from antennal lobes of 19 foragers was recorded. The experiment protocol follows previous publications with slight modifications [28]. Briefly, after a stable recording was achieved, the honeybee preparation was first stimulated with each of the following pure odors: hexanol (Hex), 2-octanone (Oct) (Sigma-Aldrich Chemie GmbH) and control (filter paper only). The duration of the pulse was 4 sec, and two minutes of recovery time were allowed between the two pulses. During the repeated exposure with Hex or Oct, 40 pulses were delivered with inter-pulse interval of 60 sec, after which 10 min recovery was given before testing with the two odors again. The order of odor presentations was randomized during the entire protocol.

### Behavioral assays and familiarization protocol

Honey bees were placed under a constant humidified air flow and vacuum system, in order to remove excess odorants. Bees were recorded from above with a web camera at 30 frames per sec as described by Birgiolas et al. (2017) [30]. The experimental protocol follows that in the electrophysiological experiments with a few differences. In brief, the honey bees were first stimulated once for 4 seconds in a randomized order with hexanol, 2-octanone, and clean air. There was a 5 min recovery period between each of the odor stimulations. Next the animals (N=24) were put through a familiarization protocol in which they received 40 pulses of either Hex (N=13) or Oct (N=11) at an inter-pulse interval of 60 sec, after which 10 min recovery was given before testing with the odors again.

### Analysis of spike data and video recordings

Extracellular spike waveforms are recorded with a TDT RZ2 system at 20 KHz sampling rate, then exported to Offline Sorter program (Plexon Inc, Dallas, TX) which classifies the similar waveforms into individual clusters (units) based on the relatedness of waveforms’ projection onto a 3D space derived from the first three principal components that capture the most variation of the original waveforms. To increase the discriminating power, the original waveforms are grouped in a tetrode configuration [57, 58], matching the physical design of the recording probe, i.e. 16 recording sites are distributed in two shanks in a block design of 2×4. Each block is called a tetrode. Statistical separation of waveform clusters, representing individual neurons or units, is aided with visual inspection, all implemented in the Offline Sorter program. Once satisfied with the clustering results, the time stamps of waveforms are then exported to Neuroexplorer program (Plexon Inc, Dallas, TX) and Matlab (Mathworks, Natick, MA) for further analysis.

Net response was calculated to quantify the stimulus triggered spike responses, using the following formula: net response = (post-stimulus response – pre-stimulus response)/pre-stimulus response. Here, the response is defined by the number of spikes within 1 or 4 sec duration after the onset of a response, which is about 0.15 sec after the opening of the solenoid valve indicated by a square pulse. After plotting out the peristimulus time histograms of all units to either Hex or Oct stimulus, it was clear that the responses have two distinct phases: transient phase and lasting phase (Fig.3 F, G). The transient phase, lasting for about 1 sec, is present for all units; but the lasting phase is less consistent, and it can be as long as the stimulus duration (4 sec) or longer or shorter. For this reason, we focused our analysis on the more consistent transient phase despite that the stimulus duration was longer.

Behavioral videos were processed in SwarmSight [30] (swarmsight.org) in order to extract positions of antenna and proboscis and were exported to R (R Core Team 2017). Statistical analyses were used to compare the antennal movements of animals in three time-windows: 4 sec immediately before, 4 sec during, and 4 sec immediately after odor presentation. Custom code in R was used to calculate the mean antennal angles for each video recording when only clean humidified air was being presented to each bee. Then for each bee and each frame included in our three time-windows (before, during, and after), a normalized angular difference from the mean antennal position was calculated. A two-way ANOVA and Tukey HSD post hoc analysis was then used to compare these normalized angular differences between neighboring time windows across animals before and after Latent Inhibition exposure to both the familiar and novel odors. We found no difference in response patterns dependent upon initial odor identity, and thus pooled odors based upon familiarity and not odor identity.

### Classification of putative PNs and LNs

We used cell-specific features to classify spike-sorted units into LNs and PNs. These features and the sorting protocol closely followed previous work [1] with one exception. We were unable to access certain cell properties (e.g., sub-threshold membrane potential) due to using extracellular recording methods. The features we used were as follows.

1. Change in firing rate. This was defined as the difference of the maximal firing rate after stimulus (within a 800ms-window) and the minimal firing rate in a 500ms pre-stimulus window. We calculated the firing rate following the method described in [1]. Briefly, for each unit, we first aligned different trials based on maximizing pairwise correlation of the time derivative [59]. We then averaged over trials and convolved this average with an asymmetric alpha kernel (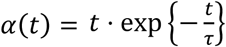, with τ = 8ms).
2. Lag to response. This was calculated as the time between the stimulus onset and the first crossing of the half of the maximal median firing rate of response, calculated over the response window.
3. Variability in the inter-spike interval. Following [1], we used a modified coefficient of variation, CV2. This metric was introduced to quantify interval dispersion when the firing rate of a cell is modulated [60]. It is defined as the variance of two consecutive inter-spike intervals divided by their mean. We then averaged this metric over all spike pairs and all trials.
4. Variability across trials. This was calculated as the Fano factor, i.e. ratio of variance to mean, of the number of spikes in the response window (800 ms) across different trials.

These four features were then used in a hierarchical clustering analysis using the Ward linkage method and the Euclidean distance metric (calculations were carried out with scipy’s *linkage* and *fcluster* functions). These features provided two well-defined clusters of distinct cell types. Projection neurons were defined by those that had larger change in firing rate due to response. Our sorted cells occupied similar regions in reduced space as the sorted data from intracellularly recorded PNs and LNs [1] (Fig. 2).

## Acknowledgement

The authors wish to thank Dr. Ozturk for maintaining the honeybee colonies. This work was funded by NIH NIGMS R01GM113967 and NSF 1556337 to BHS, DARPA HR00111990034 to BHS and MB, ONR N000141612829 to MB, and a subaward to BHS as part of the NSF/CIHR/DFG/FRQ/UKRI-MRC Next Generation Networks for Neuroscience Program.

